# Timed inhibition of CDC7 increases CRISPR-Cas9 mediated templated repair

**DOI:** 10.1101/500462

**Authors:** Beeke Wienert, Sharon J Feng, Melissa Locke, David N Nguyen, Stacia K Wyman, Katelynn R Kazane, Alexander Marson, Christopher D Richardson, Jacob E Corn

**Affiliations:** Innovative Genomics Institute, University of California, Berkeley, 94704; Department of Molecular and Cell Biology, University of California, Berkeley, CA, 94704; Gladstone Institutes, San Francisco, CA, 94158; Department of Microbiology and Immunology, University of California, San Francisco, CA, 94143; Department of Medicine, University of California, San Francisco, CA, 94143; Department of Molecular, Cellular, and Developmental Biology, University of California, Santa Barbara, California, 93106; Department of Biology, ETH Zürich, 8093 Zürich, Switzerland

## Abstract

Repair of double strand DNA breaks (DSBs) can result in gene disruption or precise gene modification via homology directed repair (HDR) from a templating donor DNA. During genome editing, altering cellular responses to DSBs may be an effective strategy to rebalance editing outcomes towards HDR and away from other repair pathways. To identify factors that regulate HDR from a double-stranded DNA donor (dsDonor), we utilized a pooled screen to define the consequences of thousands of individual gene knockdowns during Cas9-initiated HDR from a double strand plasmid donor. We find that templated dsDonor repair pathways are mostly genetically distinct from single strand donor DNA (ssDonor) repair but share aspects that include dependency upon the Fanconi Anemia (FA) pathway. We also identified several factors whose knockdown increases HDR and thus act as repressors of gene modification. Screening available small molecule inhibitors of these repressors revealed that the cell division cycle 7-related protein kinase (CDC7) inhibitor XL413 increases the efficiency of HDR by 2–3 fold in many contexts, including primary T-cells. XL413 stimulates HDR through cell cycle regulation, inducing an early S-phase cell cycle arrest that, to the best of our knowledge, is uncharacterized for Cas9-induced HDR. We anticipate that XL413 and other such rationally developed inhibitors will be useful tools for boosting the efficiency of gene modification.

## Main Text

Genome editing with targeted nucleases, such as CRISPR-Cas9, is a powerful tool for research and a promising approach for therapeutic treatment of human disease. One strategy for efficient genome editing in eukaryotic cells introduces a ribonucleotide protein (RNP) complex comprised of the type II endonuclease Cas9 and a guide RNA (gRNA), which create a double strand DNA break (DSB) at a targeted location in the genome^1,2^. This DSB is repaired by cellular DNA repair pathways to produce two outcomes: error-prone sequence disruption by insertion or deletion (indels) at the DSB, or precise sequence modification via homology directed repair (HDR) that copies a homologous donor DNA into the DSB. The targeted incorporation of introduced DNA sequences enables ground-breaking research approaches, including endogenous epitope tagging and the insertion of SNPs to test disease causation, and use of these techniques in human cells promises therapeutics to correct genetic lesions that drive human disease^3,4^. Strategies to favor precise HDR outcomes over deleterious error-prone repair in human cells are therefore of intense interest both to improve understanding of biological pathways and enable new therapeutic options.

Human cells have multiple overlapping DSB repair pathways, such as alternative-End Joining (alt-EJ), synthesis-dependent strand annealing (SDSA), and homology-directed repair, that have been implicated in Cas9-mediated gene modification^5–7^. To investigate these mechanisms in greater detail, we previously developed a reporter assay that allowed us to interrogate the genetic requirements of Cas9-mediated HDR using single stranded donor DNA (ssDonor) and discovered that single strand template repair (SSTR) requires the Fanconi Anemia (FA) DNA repair pathway^8^. We furthermore found that while HDR from a double stranded DNA plasmid (dsDonor) depends on Rad51, SSTR does not. These distinct requirements for HDR from ssDonor and dsDonor implied that different donors produce molecularly identical gene modifications via different mechanistic routes. To more completely map how different types of donors mediate Cas9-induced HDR, we used genetic screening to reveal the DNA repair factors that are involved in HDR using dsDonor DNA (a subset of HDR, from here on termed HR). Here, we describe genes that up-or down-regulate HR from a double stranded template and find pathways that are shared with and distinct from SSTR. We furthermore discover factors whose knockdown increases HDR. Timed administration of a small molecule inhibitor of one of these factors, CDC7, increases HR and SSTR by 2–3 fold in multiple types of human cells.

## Results

### SSTR and HR repair pathways are overlapping

We adapted a previously described pooled screening platform^8^ to define the contribution made by each of thousands of DNA metabolism genes to Cas9-mediated gene replacement using a plasmid donor DNA template. The basis of this platform is the stable expression of three components in each cell: 1) a dCas9-KRAB CRISPRi construct^9^, 2) a *BFP* reporter gene, and 3) a guide RNA targeting the transcription start site (TSS) of a single gene. We constructed a guide RNA library to target genes with Gene Ontology (GO) terms related to DNA metabolism, comprising a library of approximately 2,000 genes at a density of five guides per TSS^10^ **[Document S1 GUIDES]**. Pooled K562 erythroleukemia cell populations stably expressing BFP and individually inhibiting a specific DNA metabolism gene were transiently nucleofected with a Cas9 RNP targeted to introduce a DSB in the *BFP* reporter, together with a plasmid donor DNA encoding a *GFP* sequence template that will convert BFP to GFP upon successful HR^11^ **[Figure 1A]**. Edited cell populations (unedited: BFP^+^, HR: GFP^+^, gene disruption: non-fluorescent) were separated by fluorescence-activated cell sorting (FACS) and the guide RNA frequency in each population was determined by Illumina sequencing the stably integrated guide RNA cassette. Genes whose up-and down-regulation altered each repair outcome were determined by comparing the sorted populations to the starting pool. Similarities between the reagents and techniques used in this screening approach permitted direct comparison with our earlier screen editing the same locus but utilizing an ssDNA donor^8^ **[Figure 1A]**.

**Figure 1:**
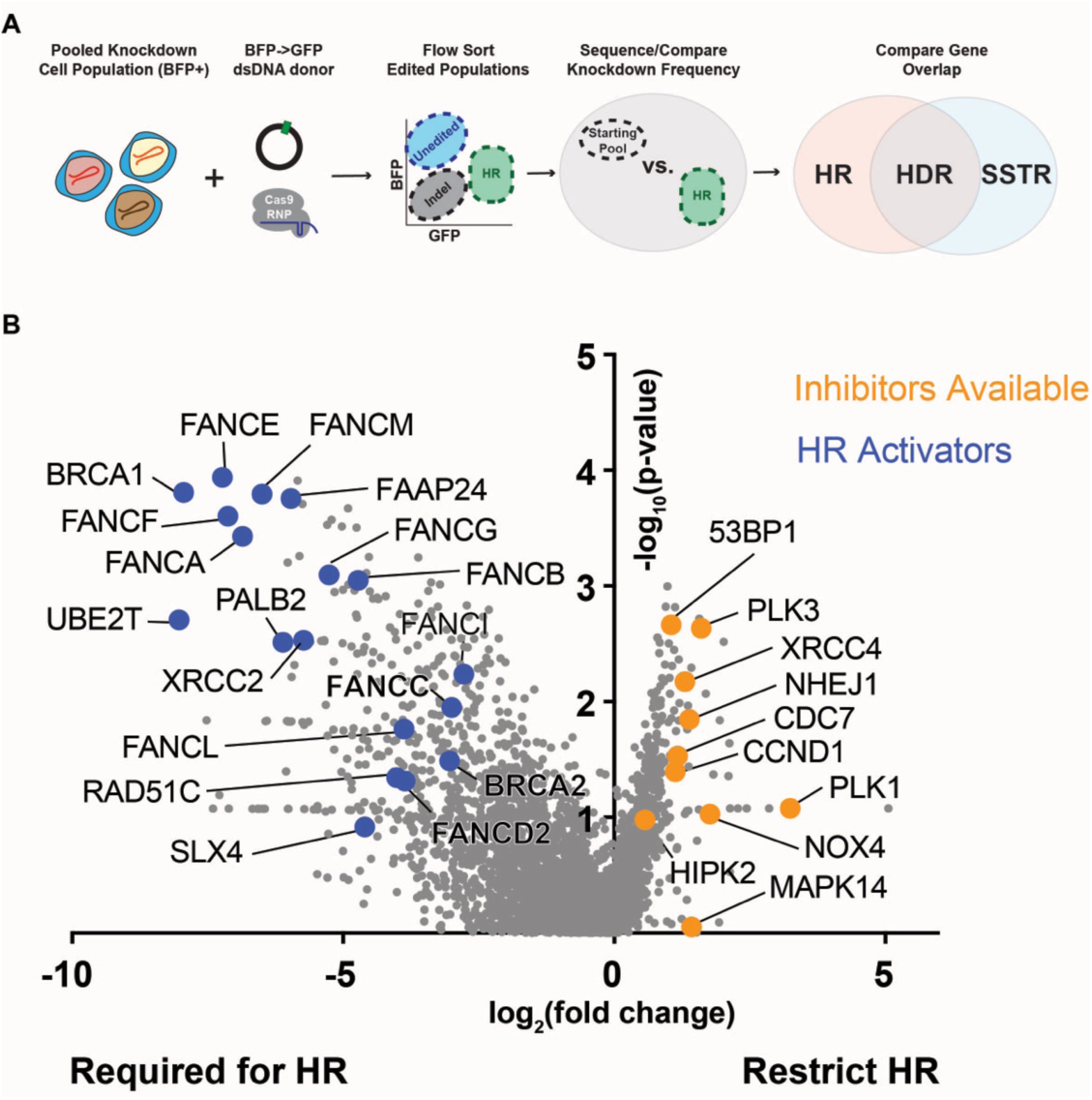
A pooled screen reveals pathways that regulate templated repair using Cas9-DSB and a double stranded plasmid donor. **(A)** Schematic showing BFP➔GFP CRISPRi screening strategy. Pooled K562 cells that stably express BFP and inhibit DNA metabolism genes are edited with Cas9 RNP that cuts within BFP and a dsDNA plasmid donor that contains a promoterless copy of GFP. Cas9 gene editing results in three populations: Unedited – BFP^+^, Indel – non fluorescent, and HR – GFP^+^. Guide RNA frequencies in the HR population were quantified and compared to the unsorted population to identify genes that promote/restrict HR. These genes were compared to data from a prior screen using ssDNA donor^8^. **(B)** Genes repress and promote dsDonor HR. A volcano plot of pooled screen hits showing the Fanconi Anemia pathway (blue) and HR repressors with available small molecule inhibitors (orange). Data presented from n=2 screen replicates.

We identified genes involved in HR by comparing guide RNA frequencies in the GFP^+^ population (*i.e*. cells that had undergone HR) to guide RNA frequencies in the unsorted control population. Guide RNAs targeting genes that restrict HR were enriched in the GFP^+^ population (because their knockdown favors HR), while guide RNAs targeting genes that are required for HR were depleted from the GFP^+^ population **[Figure 1B]**. Our dsDonor screen revealed that the Fanconi Anemia (FA) repair pathway is required for HR, which is similar to the requirement of the FA pathway for Cas9-mediated SSTR^8^. Thirty one of forty FA and FA-related genes were required for HR, suggesting that this is an activity of the overall FA pathway **[Figure 2A]**. While the FA pathway is typically associated with interstrand crosslink repair and restarting stalled replication forks, our work and those of other labs has consistently indicated that multiple FA genes play a strong role in HDR from multiple types of DNA templates^12^. While most of the FA pathway is required for both HR and SSTR, some FA factors are required for one but not the other, such as the FA E2 ubiquitin ligase, UBE2T, or the DNA binding protein, FAAP24. Moreover, components of the FA core complex including FANCA, FANCE, and FANCF showed different magnitudes of phenotypes in each screen. These results imply donor-specific functions for FA sub-complexes **[Figure 2A]**.

**Figure 2:**
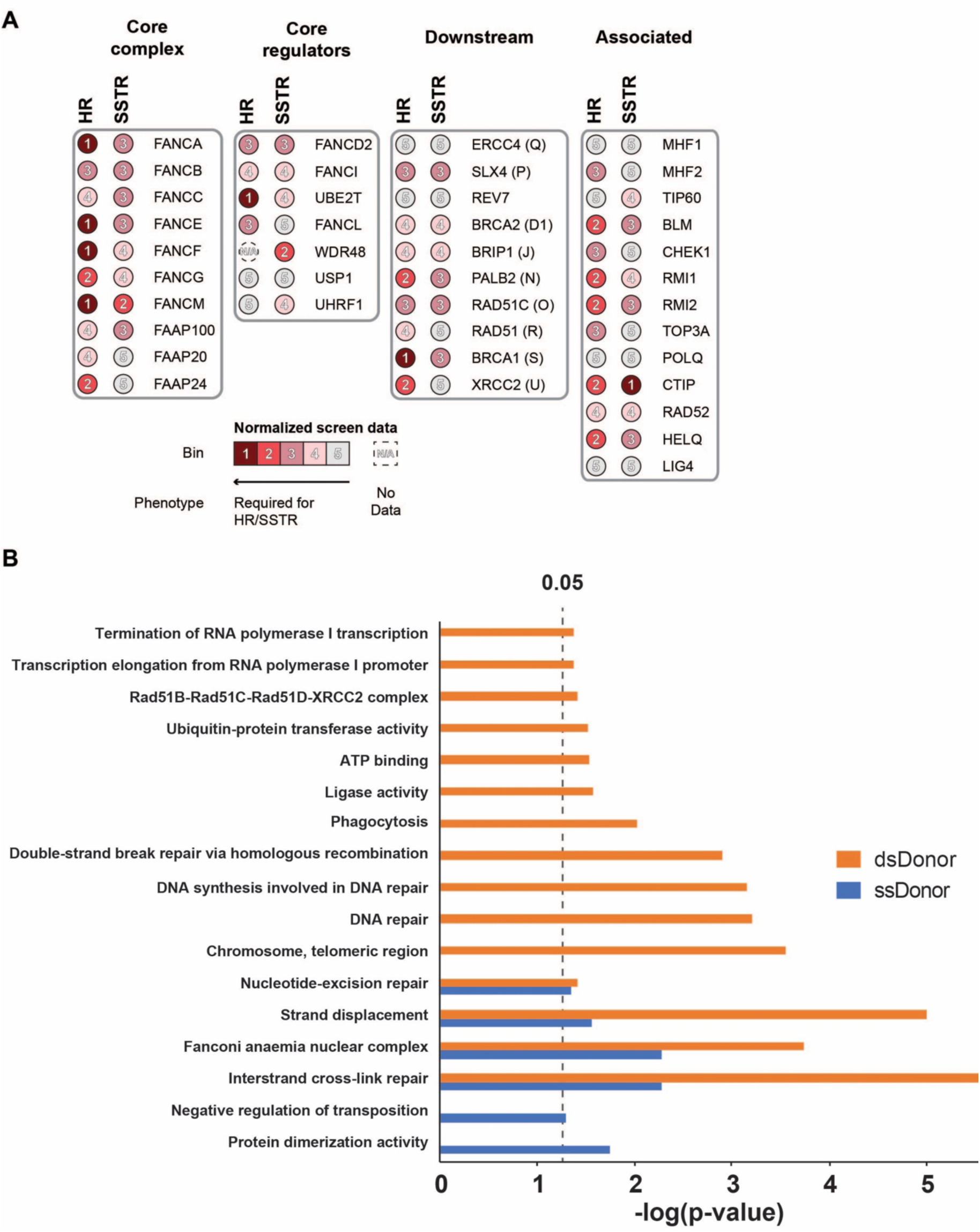
dsDonor HR requires the Fanconi Anemia pathway and is genetically distinct from SSTR. **(A)** The FA repair pathway is required for HR and SSTR. Gene names are annotated with normalized phenotype scores (see Online Methods) for both HR and SSTR screens. Increasing color intensity and decreasing bin number is proportional to the effect size seen in the HR or SSTR screen data. Functional FA complexes: the FA Core complex, Core regulators influencing FANCD2-FANCI ubiquitination, Downstream repair effectors, and Associated factors. Raw data presented in [**Document S2**]. **(B)** Unique and shared genetic pathways contribute to SSTR and HR. GO term analysis for statistically significant (p<0.05) hits from HR and SSTR screens. Data presented as GO terms enriched in HR dataset (plasmid, orange) or SSTR dataset (ssDNA, blue). Bar plot heights present statistical significance of GO term enrichment. Raw data presented in [**Document S2**] or prior publication^8^. Data represent two biological replicates each of the HR and SSTR screens.

The shared reliance of Cas9-induced SSTR and HR on the FA pathway motivated us to systematically explore overlapping genetic dependencies behind these two activities. We curated lists of statistically significant (p<0.05) genes appearing in the SSTR and HR screens and performed GO term analysis to define pathways involved in each process^13^. There was substantial overlap between SSTR and HR: both pathways require Fanconi Anemia Repair, Nucleotide Excision Repair (NER), and Strand Displacement activities, which is driven by mutual reliance on FA proteins, members of the TFIIH complex, and the BLM helicase. Shared reliance on these pathways implies that both forms of HDR may challenge cells to balance NER-like single strand editing activities and templated repair events, as has been suggested for repair of interstrand crosslinks^14^. Despite these similarities, SSTR and HR are distinct in notable ways. SSTR but not HR depends on “Negative regulation of transposition”, a GO term comprising APOBEC3C, D, F, and G. Originally reported as RNA editing enzymes, these enzymes are known to modify single stranded DNA during gene editing reactions^15^, and similar proteins have been repurposed as targeted DNA base-editing reagents^16^. Genes uniquely important for HR, on the other hand, were annotated as “Double Strand Break Repair via Homologous Recombination”, because these genes are known to play roles in well-studied forms of dsDonor-templated repair, such as meiotic homologous recombination **[Figure 2B]**. These observations suggest that HDR generally requires the FA pathway, but HR and SSTR require specialized activities to respond to donor topologies or intermediate structures specific to each repair process.

We also found several sets of genes that repress HR, and whose knockdown enhances HR efficiency **[Figure 1B]**. Some of these genes are consistent with a model in which NHEJ and HR compete to repair DSBs, and that inhibition of one pathway may favor the other^17–19^. Examples of these repressor genes include TP53-binding protein 1 (53BP1), X-ray repair cross-complementing protein 4 (XRCC4) and non-homologous end joining factor 1 (NHEJ1), which interact at DSBs to promote DNA ligase 4 (LIG4) association during non-homologous end-joining (NHEJ)^20^. Other repressors that we identified have not previously been reported to increase HR efficiency, but have roles in processes have been linked to DNA repair outcomes, such as cell cycle progression.

### Inhibiting HR repressors increases both HR and SSTR

Our exploration of genes and pathways involved in HR presented us with a number of candidate HR repressors whose knockdown increases HR efficiency. However, gene editing is frequently performed in primary cell types or in experimental contexts where transcriptional or genetic repression of these factors would be unsuitable. Small molecule treatments that increase HR would be extremely valuable because HR is quite inefficient in human cells yet desirable for its ability to precisely engineer genomic sequences and even insert long (>500 bp) sequences such as chimeric antigen receptors during T-cell engineering^21^. We performed an extensive literature search to find small molecule inhibitors of HR repressors **[Figure 1B]**. We focused on eight commercially available small molecules that are reported to inhibit CCND1, CDC7, HIPK2, MAPK14, NOX4, PLK3, PLK1, and 53BP1 **[Extended Data Figure 1A]**.

We first asked if these compounds have an effect on HR or SSTR editing outcomes using a derivative of the K562 cell line stably expressing a BFP-to-GFP reporter system^11^. Reporter cells were nucleofected with a Cas9 RNP targeting the BFP reporter gene and either an ssDNA or plasmid dsDNA donor. Following nucleofection, cells were treated with different inhibitors for 24 hours and then recovered in drug-free media **[Figure 3A]**. BFP-to-GFP HDR outcomes were monitored by flow cytometry after four days. We found that cells treated with the CDC7 inhibitor XL413^22^ showed a significant increase in both SSTR and HR **[Figure 3B]**. Inhibition of mitogen-activated protein kinase 14 (MAPK14) with SB220025 slightly enhanced SSTR and inhibition of PLK3 with GW843682X increased both SSTR and HR from a dsDonor, but the effect size was small. All other compounds resulted in no change or even a reduction of HR, which could be caused by impaired cell fitness. siRNA inhibition of CDC7 was less effective than small molecule inhibition **[Extended Data Figure 1B]** at promoting HDR, which suggests that inactivating CDC7 kinase may be more effective as an HDR stimulant than reducing the levels of CDC7 kinase. The effect of XL413 is concentration dependent, as both SSTR and HR increased in a dose-dependent manner, with 33 µM XL413 increasing HDR 2-to 3-fold **[Figure 3C]**. Importantly, XL413 concentrations up to 33 µM and exposure for up to 24h did not result in a notable decrease in viability in K562 cells **[Extended Data Figure 2A-B]**.

**Figure 3:**
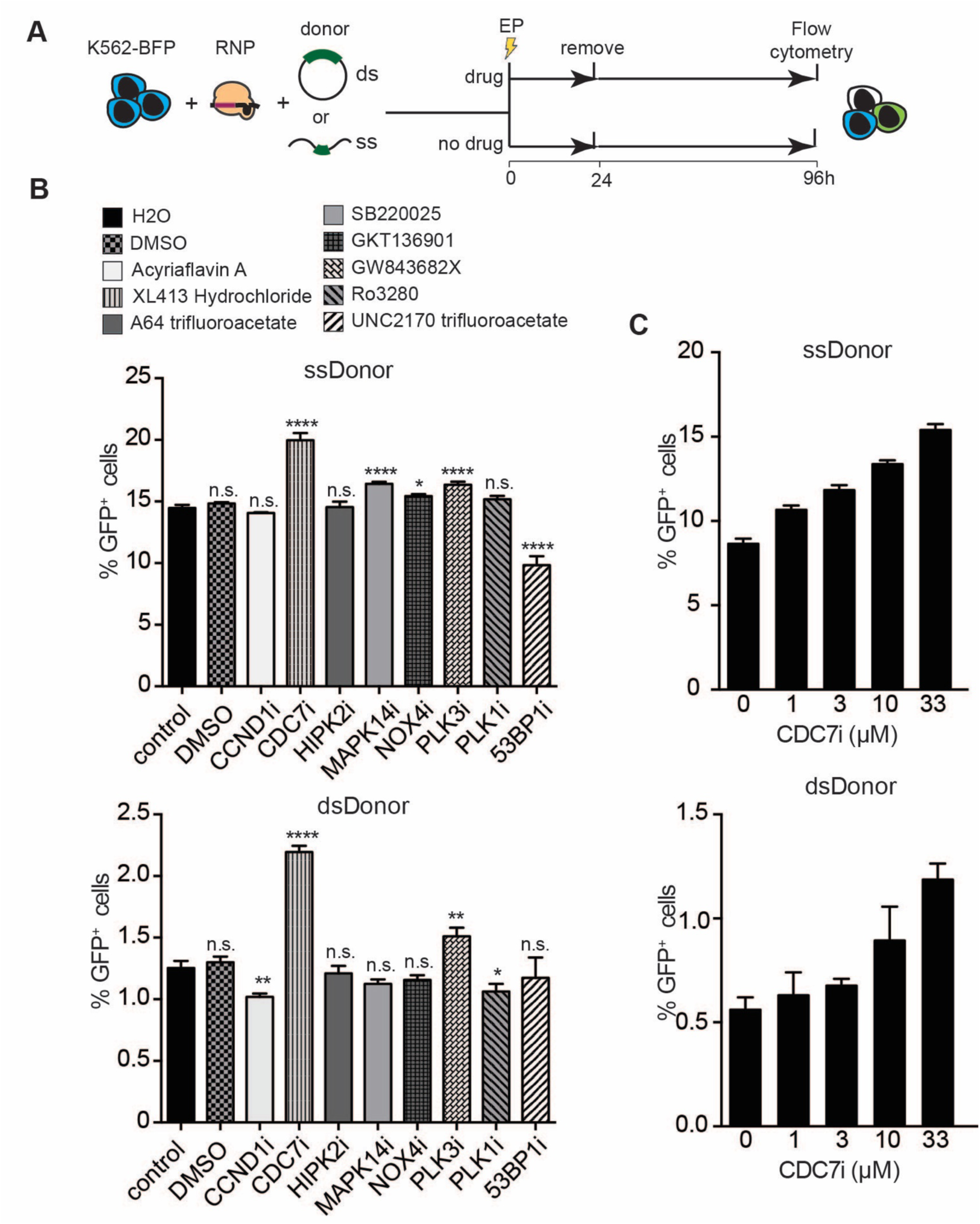
Enhancing HR by small molecule inhibition of factors discovered in genetic screening. **(A)** Schematic of small molecule evaluation. K562-BFP cells were transfected with Cas9 RNPs targeting the BFP transgene and either dsDNA plasmid or ssDNA donor. After electroporation (EP), cells were added to media with or without drug. Cell populations were recovered into fresh media after 24h and analyzed by flow cytometry after 96h. **(B)** CDC7 inhibition with XL413 significantly increases SSTR and HR by flow cytometric analysis of K562-BFP cell populations. Shown is the percentage of GFP positive cells 4 days post nucleofection with ssDNA (top) or dsDNA plasmid donor (bottom) comparing different drug treatments. X-axis indicates the intended molecular target of small molecule inhibitors, with the compound identifier above. **(C)** XL413 increases HDR in a concentration-dependent manner. Shown is the percentage of GFP positive cells 4 days post nucleofection for editing with ssDonor (top) and dsDNA plasmid donor (bottom). All values are shown as mean±SD (n=3 biological replicates). Statistical significances were calculated by unpaired t-test between indicated sample and control (*p<0.05, **p<0.01, ***p<0.001, ****p<0.0001, n.s.: not significant).

### CDC7 inhibition increases HDR at endogenous loci

We next asked if XL413’s ability to stimulate HDR is generally applicable to multiple genomic loci, cell types, and HDR “cargo” sizes. We used Cas9-induced HR to knock-in a plasmid dsDonor GFP coding sequence at the C-terminus of three different genes: *lysosomal-associated membrane protein 1 (LAMP1)*, *fibrillarin (FBL)* and *translocase of outer mitochondrial membrane 20 (TOMM20)*. These knock-in reagents were previously developed as part of a comprehensive cell-tagging effort in induced pluripotent stem cells (iPSCs)^23^. We found that treatment with XL413 for 24 hours directly after nucleofection increased the HR efficiency at *LAMP1*, *FBL* and *TOMM20* by two-to three-fold, irrespective of the original frequency of HR **[Figure 4A]**. Furthermore, we tested XL413 at the *LAMP1* locus in HCT116 and HeLa cell lines and found that these cell lines also significantly increased HR to a similar extent **[Figure 4B]**. These data demonstrate that the XL413 CDC7 inhibitor increases HR independently of the genomic locus and cell type and can be used for the installation of long sequences via HR.

**Figure 4:**
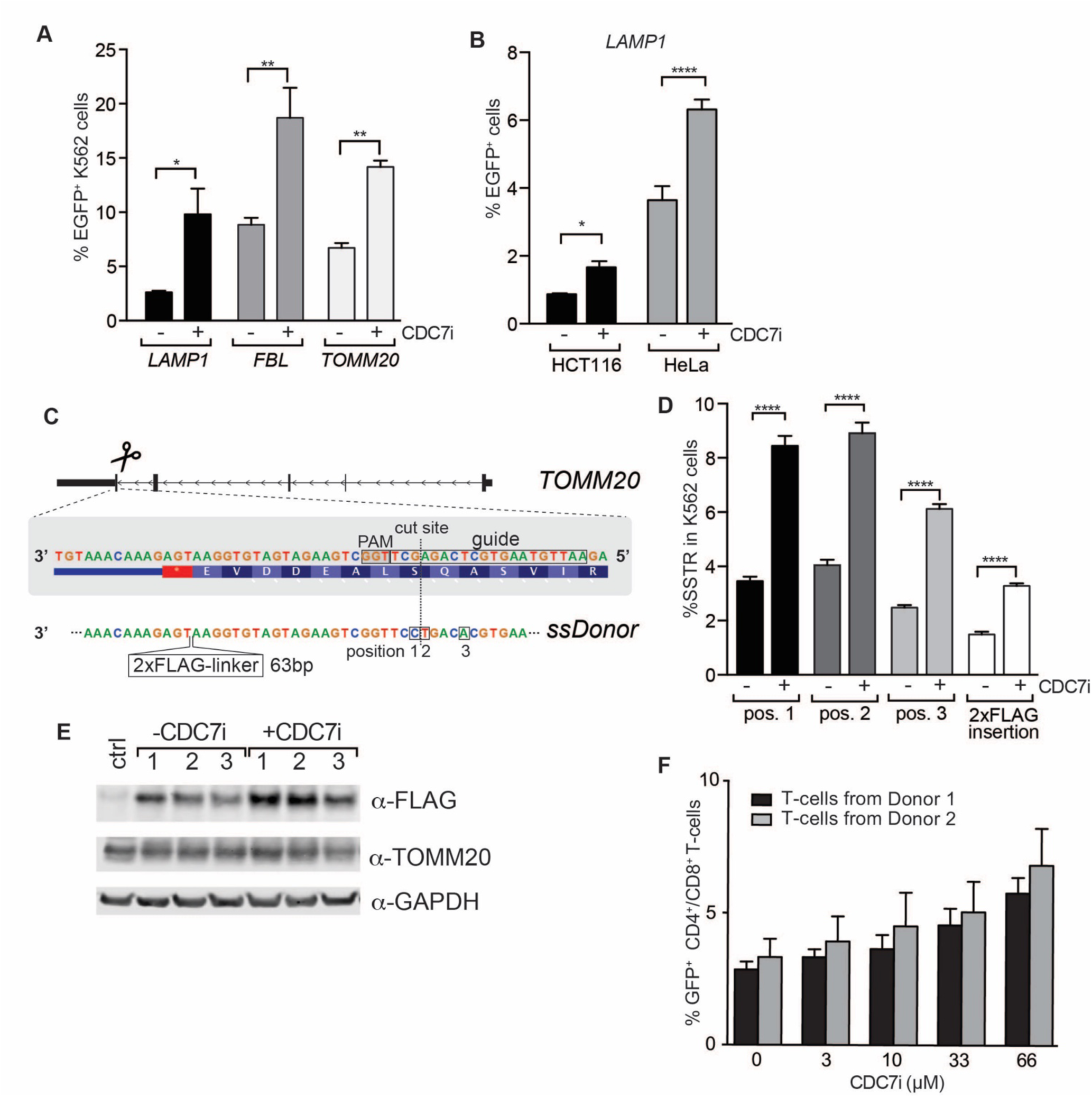
CDC7 inhibition increases HR and SSTR in diverse contexts. **(A)** XL413 increases HR at endogenous loci with large HR donors. Cell were nucleofected with RNP targeting three different genomic loci *(LAMP1, FBL* and *TOMM20)* and dsDNA plasmid donor DNA to knock-in an *eGFP* sequence to the C-terminal end of the gene. Half of the pool of nucleofected cells was treated with 33 μM XL413 for 24h while the other half remained untreated. Flow cytometric analysis determined the percentage of eGFP positive K562 cells 4 days after nucleofection. **(B)** XL413 increases HR in multiple cell lines. Flow cytometric analysis of HCT116 and HeLa cells edited at the *LAMP1* using the same editing strategy as described in **(A)**. **(C)** Schematic outlining the genome editing strategy to knock-in a 2xFLAG sequence at the C-terminus of using an ssDonor. A linker (GGGGS)-2xFLAG sequence (63 bp) and three additional silent point mutations are introduced near the Cas9 target site. **(D)** XL413 increases SSTR at endogenous loci. Cells were nucleofected with RNP targeting *TOMM20* and 2xFLAG ssDonor [Figure 4C] in the presence or absence of 33 μM XL413 for 24h, gDNA was extracted after 4 days, and SSTR frequencies were determined by amplicon sequencing. **(E)** XL413 promotes on-target editing. Western Blot analysis for FLAG and TOMM20 expression in non-transfected K562 cells (ctrl) and nucleofected K562s cells with and without treatment with XL413. Cells were nucleofected with RNP targeting *TOMM20* and 2xFLAG ssDonor in the presence or absence of 33 μM XL413 for 24h. Cells were harvested for protein extraction 4 days post nucleofection. Data presented is representative of Western Blots on n=2 experiments. **(F)** XL413 increases HR efficiency in primary human T-cells. CD3+ T-cells (mixture of CD4+ and CD8+) from two healthy donors were nucleofected with RNP targeting *RAB11A* and 0.25 μg of a linear dsDNA donor that encodes an N-terminal fusion of GFP to the *RAB11A* gene. XL413 was added to growth media for 24h post-editing in indicated concentrations, and GFP expression was determined by flow cytometry after 3 days (n=3 per condition for each donor). Values in panels A-F are shown as mean±SD (n=3 biological replicates). Statistical significances were calculated by unpaired t-test (*p<0.05, **p<0.01, ***p<0.001, ****p<0.0001, n.s.: not significant).

We investigated if SSTR is similarly increased by CDC7 inhibition at an endogenous locus. We designed an editing strategy that uses an ssDonor to insert a 2xFLAG tag and linker at the C-terminus of *TOMM20* **[Figure 4C]**. To avoid re-cutting the repaired locus, we introduced three additional silent substitutions into the donor template to remove the gRNA recognition site. A hallmark of SSTR is a strong decrease in donor sequence incorporation with increasing distance from the Cas9-cut site^24^. However, sometimes PAM sites are unavailable at the exact introduction site reducing knock-in efficiencies dramatically. Increasing SSTR efficiency in such a context would be particularly helpful. We incorporated this scenario into our experiments by designing the ssDonor so that the tag insertion is 20 bp from the Cas9 cut site. Using amplicon PCR and next-generation sequencing, we found that XL413-treated cells again had a two-to three-fold increase in SSTR relative to untreated cells, significantly boosting the insertion of the FLAG-tag, despite its distance from the Cas9-cut site **[Figure 4D]**. Sequence-level increases in tag insertion corresponded to increased ability to detect FLAG-tagged TOMM20 by Western blotting **[Figure 4E]**. These findings suggest that CDC7 inhibition robustly increases SSTR and HR, and in this context can be used to increase the frequency of both single nucleotide substitutions and endogenous gene tagging.

### CDC7 inhibition enhances knock-in efficiency in primary T-cells

HDR in primary cells is a long-standing goal of gene editing, both for its ability to correct disease-causing SNPs and to deliver large payloads such as chimeric antigen receptors^21,25^. We therefore investigated the ability of XL413 to increase HR in human T-cells derived from healthy donor peripheral blood mononuclear cells (PBMCs). We performed editing using an RNP targeting the *RAB11A* locus and a linear dsDNA donor to generate an N-terminal GFP fusion^21^. XL413 treatment after editing produced a dose-dependent increase in HR efficiency that approached two-fold over the untreated control **[Figure 4F]**, without evidence of decreased viability **[Extended Data Figure 3]**. XL413’s ability to potentiate Cas9-mediated HR may make it valuable for challenging gene editing workflows in T-cells and other primary cell types, for example when making CAR-Ts. We speculate that XL413 or alternative CDC7 small molecule inhibitors could improve the overall fraction of successfully edited cells in these contexts.

### CDC7 inhibition causes cell cycle arrest

We next examined the molecular mechanism by which the XL413 CDC7 inhibitor leads to increased HDR. CDC7 is the catalytically active subunit of DBF4-dependent kinase (DDK), which phosphorylates and activates the MCM helicase to initiate the G1/S transition^26,27^. We therefore expected XL413 to halt cells in G1/S instead of late S/G2 when HR is supposed to be most active. We found that XL413 treatment indeed rapidly and reversibly inhibited MCM2 phosphorylation **[Figure 5A]**, but that XL413 treatment caused accumulation of cells in S-G2-M phases of the cell cycle, as measured by a FUCCI live cell cycle reporter and propidium iodide staining^28^ **[Extended Data Figure 4 and Figure 5B]**. Arrest with XL413 is distinct in cell cycle from other cell cycle modulators that have been tested for increased HR such as Aphidicolin and Hydroxyurea ^7^ **[Figure 5C-D]**.

**Figure 5:**
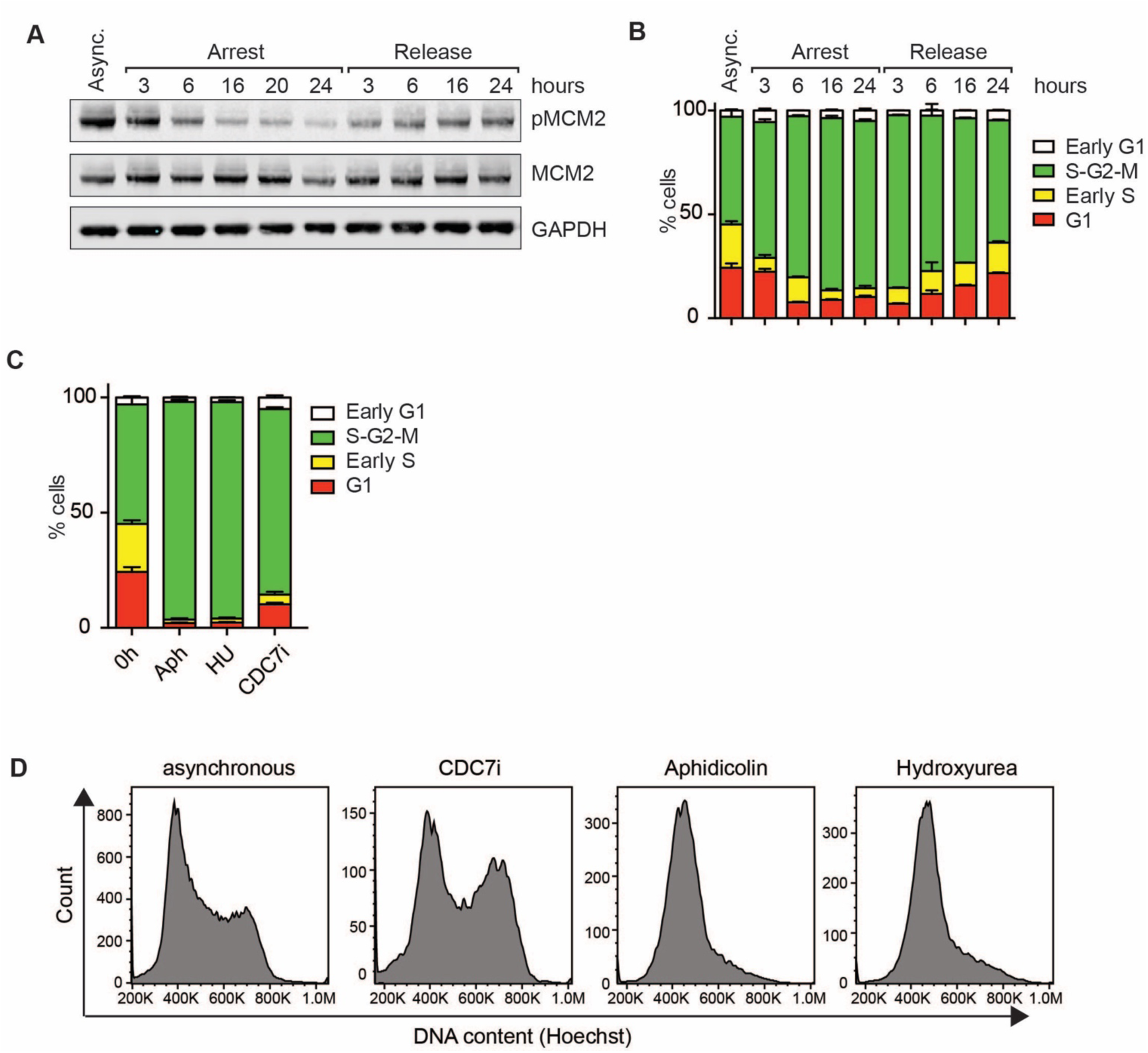
CDC7 inhibition promotes reversible cell cycle arrest. **(A)** CDC7 inhibition reduces phospho-MCM2 by Western Blot. K562 cells were either left asynchronous or treated with XL413. Cells were harvested for protein extraction at indicated time points during arrest-release experiments. **(B)** XL413 induces a reversible cell cycle arrest. Representative flow cytometry plots of asynchronous K562-FUCCI cells and K562-FUCCI cells arrested with XL413 for the indicated time. **(C)** CDC7 inhibition concentrates cells in S-G2-M. Cell cycle status in asynchronous K562 FUCCI^28^ cells and K562 FUCCI cells treated with XL413. Cells were treated as described in panel **(A)** and analyzed by flow cytometry. **(D)** CDC7 inhibition is distinct from arrests produced by Aphidicolin or Hydroxyurea, as measured by propidium iodide staining for DNA content. Cells were arrested with XL413 (33 μM), Aphidicolin (2 μg/mL), or Hydroxyurea (2 mM) for 24h. Cells were harvested 24h after arrest, stained with Hoechst 33342 to monitor DNA content, and analyzed by flow cytometry. Data presented as in panel **(C)**. Data in all panels are representative of three independent experiments.

### Timing of CDC7 inhibition determines its efficacy

Since CDC7 is an initiator of the G1/S transition, we reasoned that even a slight alteration of the timing of XL413 administration should dramatically alter cell cycle distribution. We previously arrested cells using a timing that would cause them to accumulate in G1/S *during* editing by “post” exposure to CDC7 inhibitor (e.g. edited cells recovered into XL413-containing medium) **[Figure 4]**. We therefore asked whether accumulating cells in G1/S and then *releasing* them during editing by “pre” exposure to XL413 would move them from an HR permissive to non-permissive section of the cell cycle.

Our usual post-exposure to XL413 after RNP and donor nucleofection supported increased levels of SSTR and HR **[Figure 6A]**. However, pre-exposure to XL413 and then release during editing resulted in reduced levels of HDR, suggesting that timing of cell cycle arrest and releasing the cells into HDR-permissive S-phase is crucial. The cell cycle arrest caused by XL413 presumably changes three major parameters in the cell: the activity of CDC7, the frequency of replication fork-DSB encounters, and the activation of cell-cycle regulated DNA repair pathways^29^. While Aphidicolin and Hydroxyurea also arrested cells at the G1/S transition, these arrests preserved a G1 DNA content **[Figure 5C-D]**, and did not support increased HR across multiple sites when used with the same drug-administration scheme as XL413 (i.e. edit, recover into drug, then transfer to drug-free media) **[Figure 6B]**. Together with our pre- vs post-inhibition experiments, this suggests that early-to-mid S-phase is a critical window for Cas9-mediated SSTR and HR, rather than the assumed late S/G2.

**Figure 6:**
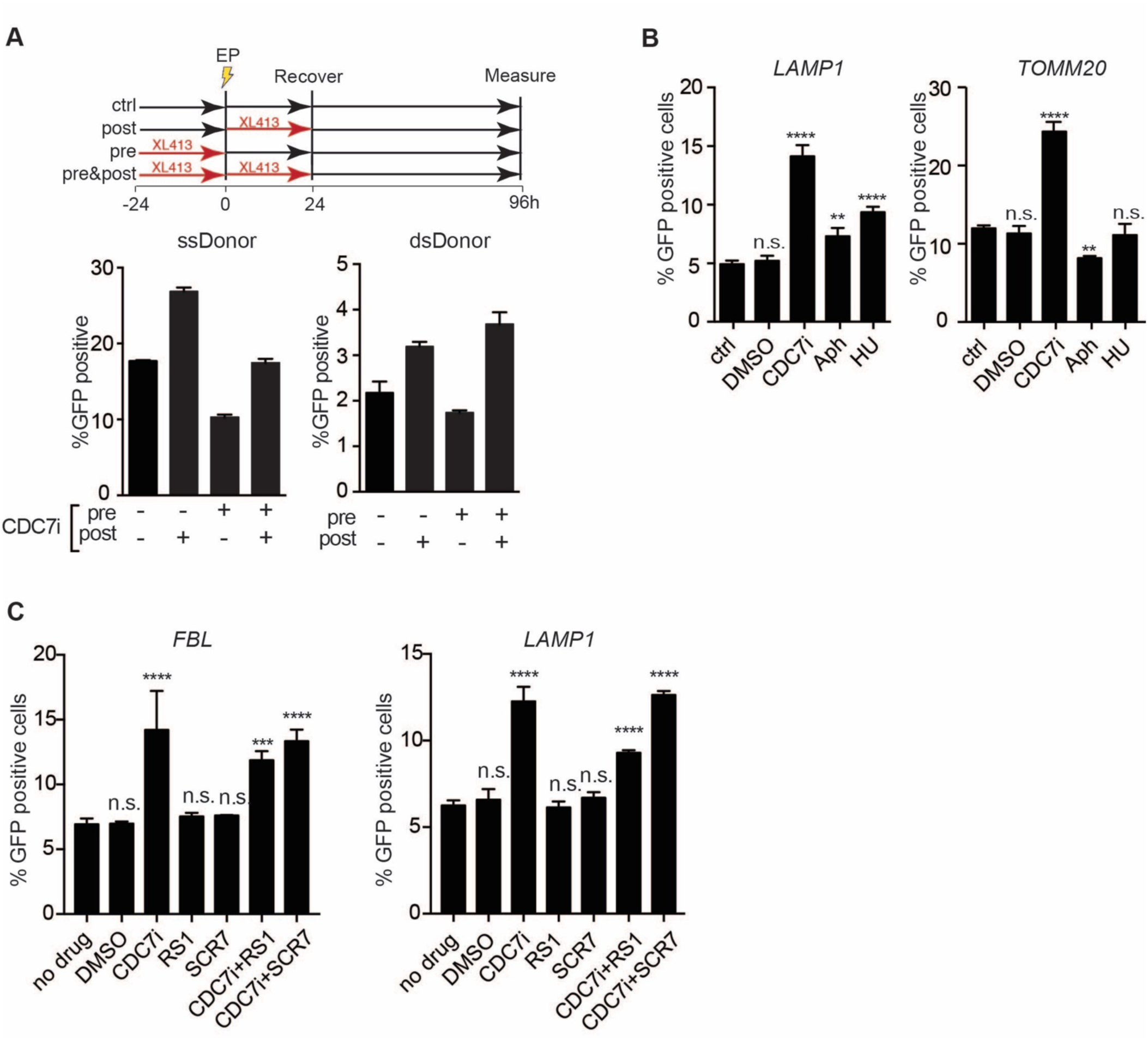
The timing of cell cycle arrest and release by CDC7 inhibition during editing is critical for SSTR and HR. **(A)** Editing outcomes in K562-BFP cells after nucleofection with RNP and donor DNA targeting the BFP transgene. Cells were untreated, treated for 24h with XL413 before nucleofection (pre), treated for 24h with XL413 after nucleofection (post) or both (pre- and post). Percentage of eGFP positive cells was determined 4 days post nucleofection using flow cytometry. **(B)** CDC7-induced arrest is more effective at boosting HR than other cell cycle arrests. K562 cells were nucleofected with RNP and eGFP donor plasmids targeting the C-terminus of either *LAMP1* (left panel) or *TOMM20* (right panel). Small molecules were added to the media of nucleofected cells for 24h. eGFP expression was monitored by flow cytometry 4 days post-nucleofection. **(C)** CDC7 arrest does not synergize with other HR boosting treatments. Effect of XL413 (33 μM), SCR7 (1 μM) and RS-1 (10 μM) treatment on editing outcomes in K562 cells. Cells were nucleofected with RNPs and eGFP plasmid donor DNA targeting *FBL* and *LAMP1* loci and small molecules were added to the media of nucleofected cells for 24h. Cells were analyzed by flow cytometry 4 days post nucleofection. All values are shown as mean±SD (n=3 biological replicates). Statistical significances were calculated by unpaired t-test (*p<0.05, **p<0.01, ***p<0.001, ****p<0.0001, n.s.: not significant).

Finally, we asked how CDC7 inhibition by XL413 compared to other small molecule approaches that have been reported to boost HR and SSTR. The LIG4 inhibitor SCR7 reportedly increases HR by inhibiting NHEJ, and the RAD51 agonist RS-1 reportedly increases HR by boosting recombination itself^30,31^. We tested both SCR7 and RS-1 for their ability to increase HR and SSTR at two different endogenous loci but found no stimulation of either type of HDR. Combining SCR7 or RS-1 with XL413 for 24 hours similarly did not increase HR beyond XL413 alone, and RS-1 treatment may actually reduce CDC7-inhibition effects on HR **[Figure 6C]**.

## Discussion

In summary, we have defined a network of genes that contribute to Cas9-mediated HR using dsDonor DNA. A close comparison between the results from two pooled CRISPRi screens using different donor templates (ssDNA vs. plasmid dsDonor) revealed that many DNA repair factors are shared between SSTR and HR. The most striking commonality is the shared dependence on much of the FA pathway for both forms of repair. Despite this shared reliance on the FA pathway, we also observed genetic differences between the different types of HDR as well as more subtle differences in the requirement for certain components of the FA subcomplexes. Overall, this suggests that the fundamental HDR pathway is the same for templated repair of a Cas9 break with ssDonor or dsDonor, but the stability and incorporation of different donor templates requires different factors. Future work could dissect the roles of the different FA sub-complexes in SSTR and HR.

We mined our HR and SSTR screening datasets to identify factors that repress HDR and whose knockdown increases the efficiency of targeted genome modifications, such as the introduction of point mutations or knock-in of fluorescent reporters. We used this genetic knowledge to develop effective protocols to enhance both SSTR and HR through small molecule inhibition of CDC7. XL413, a CDC7 inhibitor, improves the efficiency of gene editing workflows for basic research or therapeutic applications, dramatically increasing gene replacement in challenging contexts, in multiple cell types, and in cell types with tremendous therapeutic potential, such as primary T-cells. This increase in gene replacement efficiency is caused by cell cycle arrest and is consistent with existing models that HR pathways are mainly active during S/G2/M phases^29^. However, our results suggest that the HDR-permissive window may be much narrower than previously supposed. In our hands, arrest of cells in G1/S with the DNA polymerase inhibitor Aphidicolin, or the ribonucleotide reductase inhibitor Hydroxyurea, does not increase HDR. Nocodazole arrest of cells at G2/M prior to nucleofection has been reported to increase HDR in cell lines. However, reduction of this stimulation when Nocodazole-arrested cells are released into an Aphidicolin block suggests that release from Nocodazole into a *subsequent* S-phase is the key parameter, and not the G2/M arrest itself^7^. Collectively, these data suggest that XL413 increases the percent of cells in early S phase, when Cyclin-dependent Kinases are active and DNA repair protein abundance is high, and that these parameters in turn support increased levels of HDR.

Our identification of XL413 through systematic characterization of DNA repair pathways demonstrates the robustness of our approach. XL413 works best when administered after Cas9 editing, despite the fact that CDC7 inhibition was identified in a screen that was performed in the context of constitutive transcriptional inhibition (both pre-and post-editing) of DNA repair factors. These methodological differences between our screening platform and downstream small molecule inhibition may explain why small molecule inhibition of many screen hits was suboptimal. Stronger methodological ties between screen and validation may thus be the basis for future screening approaches using these reagents.

We speculate that additional combinatorial treatments could be used to boost HDR beyond what can be achieved with CDC7 inhibition alone. In our hands, the two previously described small molecules RS-1 and SCR7^30,31^ did not have an additive effect with XL413 to increase HR. However, we have not exhaustively tested XL413 in combination with other small molecules that increase HR, such as Beta adrenergic receptor agonists^32^. Nor have we tested co-administration of recombinant proteins, such as the 53BP1 inhibitor reported to boost HDR outcomes in human cell lines^33^. Our unbiased screen also identified that knockdown of 53BP1 increases HDR, so dual inhibition of CDC7 and 53BP1 could further increase HDR. However, 53BP1 cooperates with p53 to suppress genomic instability^34^, and it is unclear what effect misregulating this protein to boost HDR may have on genome stability.

The recent development of cell-cycle regulated Cas9 derivatives, including a Cas9-Geminin fusion, introduces the possibility of restricting Cas9 activity to HDR-permissive S/G2/M phases and is complementary to manipulation of the cell cycle^35,36^. While use of this Cas9 variant marginally increases absolute HDR efficiency, it decreased unwanted NHEJ since Cas9-Geminin is degraded in G1 phase when NHEJ is the main repair pathway. Combining cell-cycle regulation of Cas9 activity with accumulation of cells in the permissive phase of the cell cycle may thus comprise a potent strategy to boost HDR while minimizing undesirable editing outcomes. We anticipate that further work to map fundamental DNA repair pathways will suggest new strategies and targetable regulators to increase the precision and the efficacy of gene editing workflows.

## Acknowledgments

J.E.C is supported by a National Institutes of Health New Innovator Award (DP2 HL141006), the Li Ka Shing Foundation, and the Heritage Medical Research Institute. A.M. received a career award from the Burroughs Wellcome Fund, funding from Innovative Genomics Institute (IGI), and is a Chan Zuckerberg Biohub investigator. B.W. was funded by a Sir Keith Murdoch fellowship from the American Australian Association. C.D.R. was funded by the Fanconi Anemia Research Fund.

## Author Contributions

B.W., J.E.C., and C.D.R. conceived the study. C.D.R. and B.W. designed experiments. S.J.F. and K.R.K. performed pooled CRISPRi screen. C.D.R. analyzed data from CRISPRi screen. M.L. generated K562 FUCCI reporter cell line. D.N. performed T-cell experiments. B.W. and S.J.F. performed drug treatments and genome editing in all cell lines. B.W. performed cell cycle experiments with FUCCI cells. S.K.W. performed NGS data analysis. A.M. provided reagents. C.D.R. and J.E.C. supervised study. C.D.R., B.W., S.J.F. and J.E.C. wrote the manuscript with input from all authors.

## Supplemental Documents

Document S1 -All molecular biology reagents

Document S2 -Pooled screen results

## Extended Data Figures

**Extended Data Figure 1:**
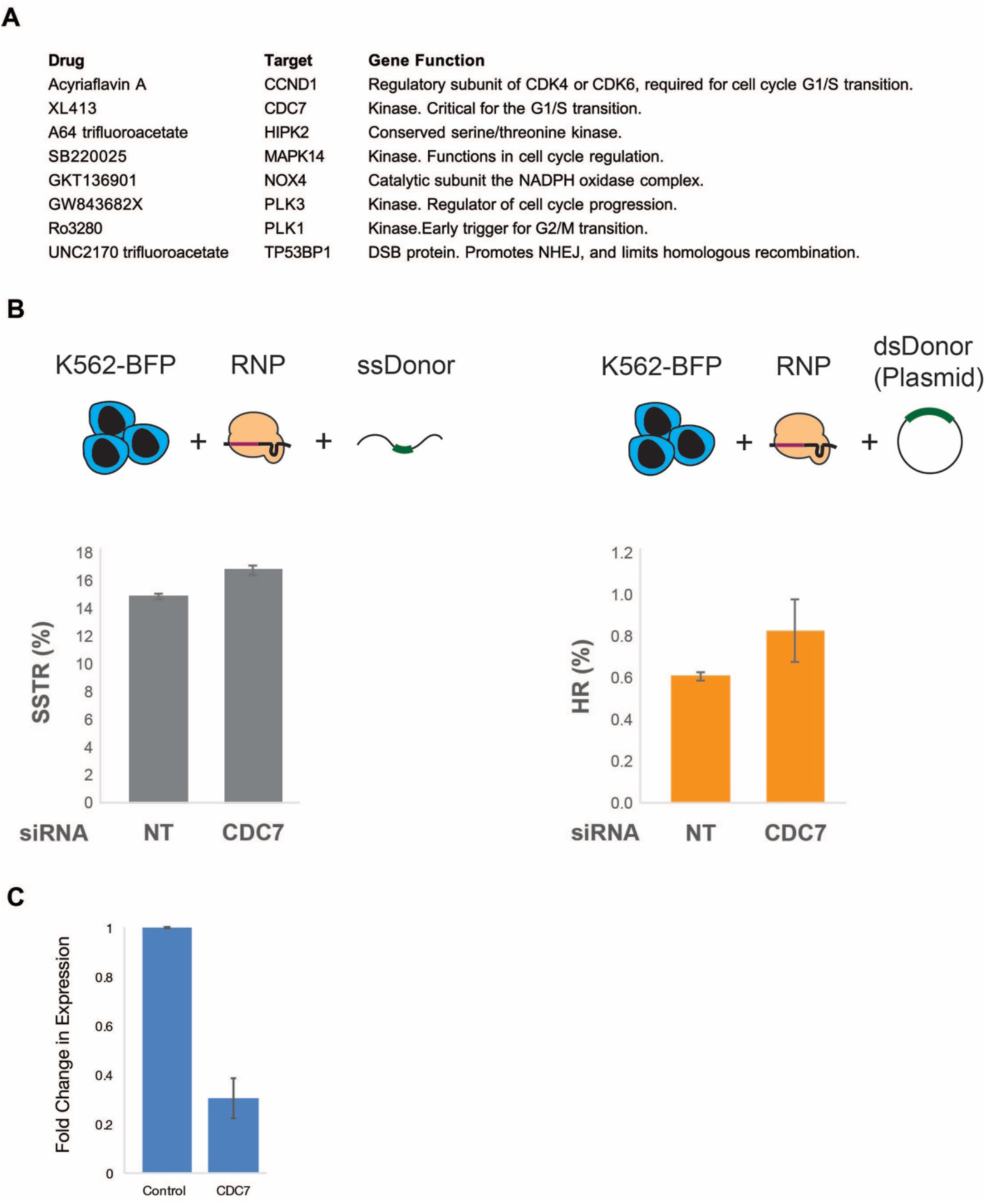
siRNA inhibition of factors that restrict SSTR, HR, or both can promote templated repair events. **(A)** Factors restricting SSTR and HR can themselves be inhibited by small molecules. HDR repressors identified from our screen [**Figure 1**] that can be targeted with known small molecule inhibitors. **(B)** siRNA inhibition of CDC7 influences SSTR and HR. Cells treated with siRNA against mock sequence (NT) or CDC7 were edited with the indicated RNP and donor DNA and %GFP was plotted for each population as analyzed by flow cytometry. Data presented as mean±SD (n=2). **(C)** Fold depletion of the target transcript over controls (ACTB, GAPDH) was measured by qPCR. Data presented were calculated from n=4 cell pellets harvested at the time of electroporation.

**Extended Data Figure 2:**
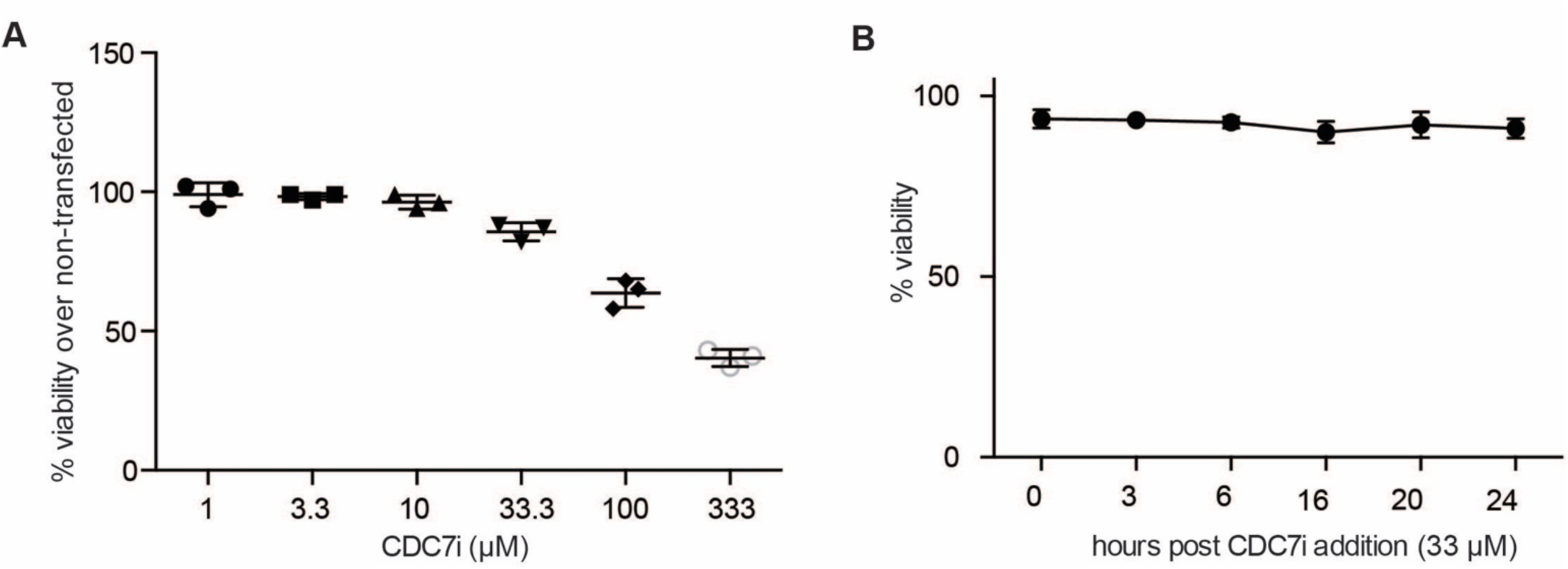
Effect of XL413 inhibitor on viability of K562 cells. **(A)** Increasing doses of XL413 are toxic. Viability of K562-BFP cells after nucleofection and treatment with XL413 at indicated concentrations for 24h. Viability was determined after 4 days by flow cytometry using forward and side scatter gates and viability was normalized to non-transfected K562 cells. **(B)** Working dose of XL413 is well tolerated. Viability of K562 cells at different time points post XL413 addition (33 μM). Viability was determined using Trypan blue exclusion test.

**Extended Data Figure 3:**
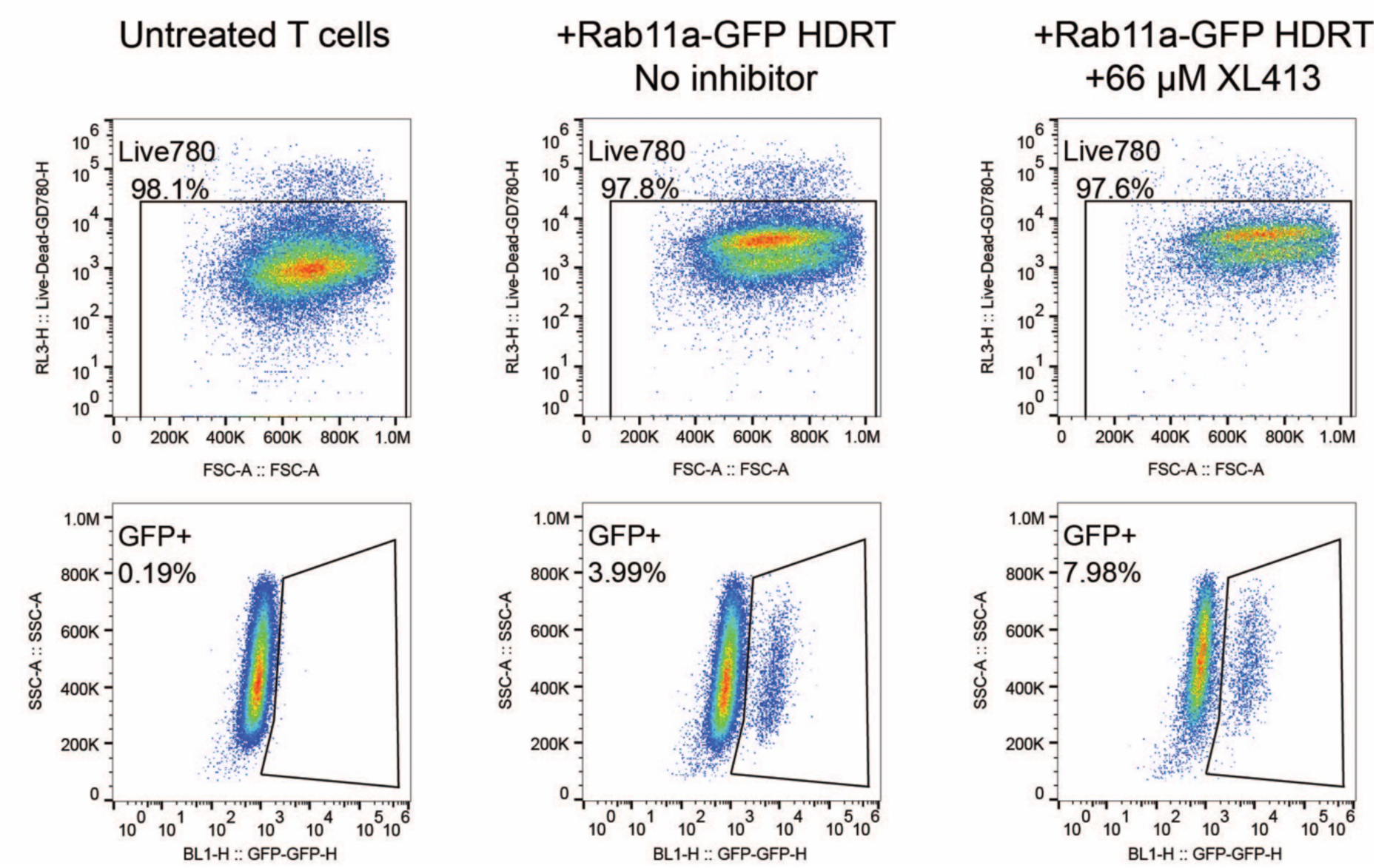
Effect of XL413 inhibitor on growth of primary human T cells. CD3+ T-cells (mixture of CD4+ and CD8+) from two healthy donors were nucleofected with RNPs targeting *RAB11A* and 0.25 μg of a linear dsDonor template DNA that encodes an N-terminal fusion of GFP to the *RAB11A* gene. XL413 was added to a growth media for 24h post-editing in indicated concentrations (n=3 per condition for each donor). Viability was determined by staining with GhostDye780, and cell count was determined by sampling equal volumes per well on an Attune NxT flow cytometer. Panels from representative samples showing flow cytometry plots depicting viability (top) and GFP positivity (bottom) from electroporated CD3+ T cells at day 3 that are either not treated with XL413, nucleofected with RNP and RAB11A-GFP donor but not treated with XL413, or nucleofected with RNP and RAB11A-GFP donor and treated with 66 μM XL413.

**Extended Data Figure 4:**
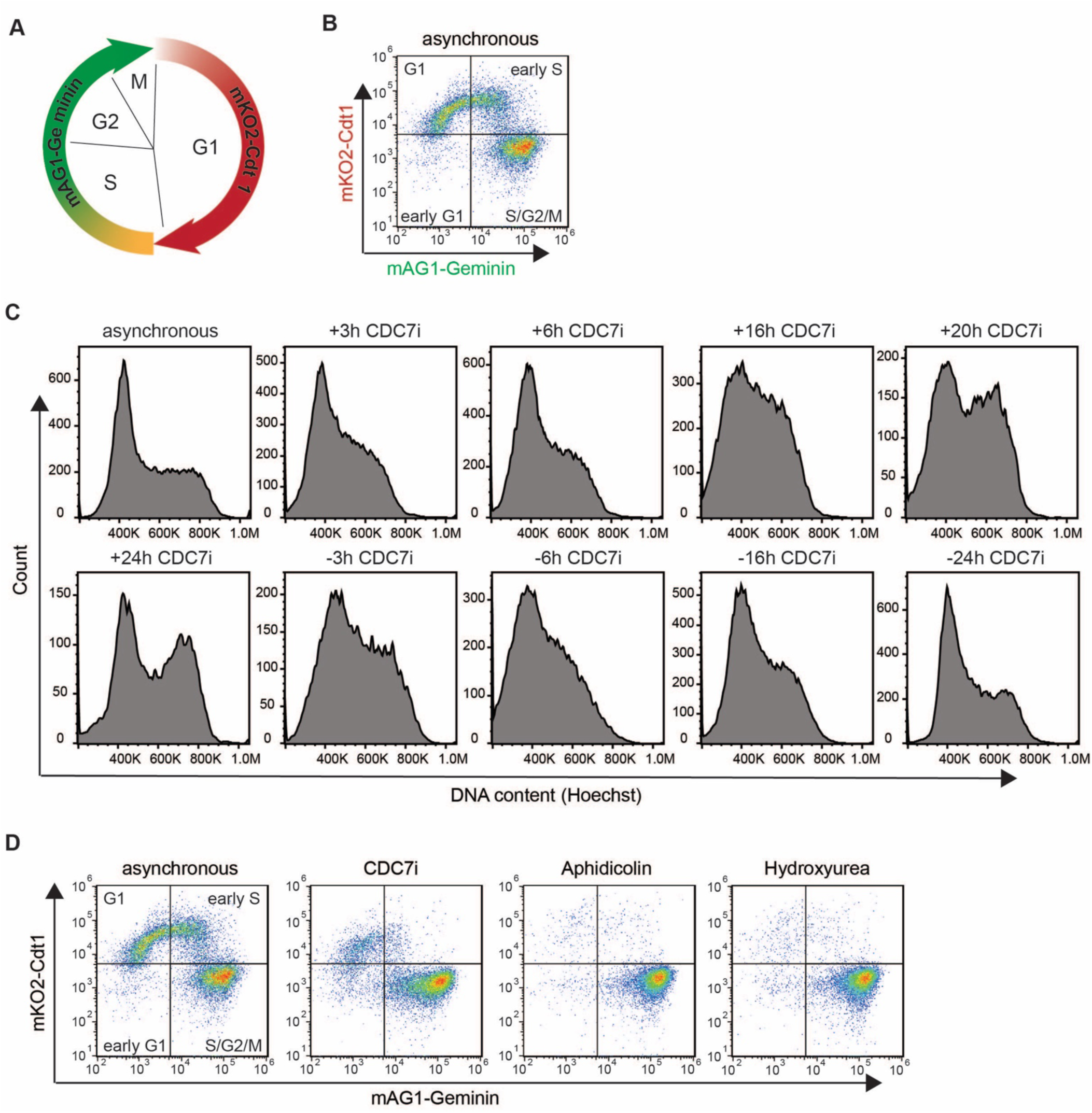
CDC7 inhibition promotes reversible cell cycle arrest. **(A)** Schematic showing the cell cycle dynamics of the FUCCI system. **(B)** Flow cytometric analysis of asynchronous K562-FUCCI cells. **(C)** CDC7 arrest reversibly alters DNA content of cell line cultures. DNA content of K562-FUCCI cells that were treated with XL413 (33 μM) for 24h and then released into media without XL413. Cells were stained with Hoechst33342 before being subjected to flow cytometry at the indicated timepoints. **(D)** Flow cytometric analysis of arrested K562-FUCCI cells.

## Online Methods

### Cell Lines and Culture

HEK293T, HCT116, HeLa and K562 cells were acquired from the UC Berkeley Tissue Culture Facility. HEK293T, HCT116 and HeLa cells were maintained in DMEM medium supplemented with 10% fetal bovine serum and Penicillin/Streptomycin. K562 cells were maintained in RPMI medium supplemented with 10% fetal bovine serum and Penicillin/Streptomycin. Cell lines were tested regularly for mycoplasma contamination using enzymatic (Lonza, Basel, Switzerland) and PCR-based assays (Bulldog Bio, Portsmouth, New Hampshire).

### Cas9, RNA, and Donor DNA Preparation

Streptococcus pyogenes Cas9 (pMJ915, Addgene #69090) with two nuclear localization signal peptides and an HA tag at the C-terminus was expressed in Rosetta2 DE3 (UC Berkeley Marcolab) cells. Cell pellets were sonicated, clarified, Ni2+ -affinity purified (HisTraps, GE life sciences), TEV cleaved, cation-exhanged (HiTrap SP HP, GE life sciences), size excluded (Sephacryl S-200, GE life sciences) and eluted at 40 mM in 20 mM HEPES KOH pH 7.5, 5% glycerol, 150 mM KCl, 1 mM dithiothreitol (DTT) ^37^.

sgRNAs were synthesized by Synthego as modified gRNAs with 2’-O-methyl analogs and 3’ phosphorothioate internucleotide linkages at the first three 5’ and 3’ terminal RNA residues using protospacer sequences described in [**Document S1**].

cRNAs/TracrRNAs were chemically synthesized (Edit-R, Dharmacon Horizon) using protospacer sequences described in [**Document S1**].

ssDonor was obtained by ordering unmodified Ultramer oligonucleotides (Integrated DNA Technologies). dsDonor was obtained by purifying plasmid DNA from bacterial cultures containing the indicated plasmid (Qiagen) or by SPRI purification of long double-stranded PCR amplicon.

### Plasmid Constructs

Sequences for plasmids used in this study are described in [**Document S1**]. Plasmids and maps will be available on Addgene after publication.

### Cas9 RNP Assembly and Nucleofection

30 pmoles of sgRNA was diluted using Cas9 buffer (20 nM HEPES [pH 7.5]), 150 mM KCl, 1mM MgCl_2_, 10% glycerol, and 1 mM TCEP). 0.75 μl of 40 μM Cas9-2xNLS (30 pmoles) was slowly mixed in, and the resulting mixture was incubated for five minutes at room temperature to allow for RNP formation. After incubation, either 0.3 μl of 100 μM ssDonor or 2 μg of plasmid DNA was introduced and mixed by pipetting. The total volume of RNP solution was 5 μl, where the volume of Cas9 buffer was adjusted to account for volume differences between ssDonor and plasmid DNA. Between 1e+05 and 2e+05 cells were harvested, washed once in PBS, and resuspended in 15ul of nucleofection buffer (Lonza, Basel, Switzerland). 5ul of RNP mixture was added to 15ul of cell suspensions. Reaction mixtures were electroporated in Lonza 4D nucleocuvettes, incubated in the nucleocuvette at room temperature for five minutes, and transferred to culture dishes containing pre-warmed media. Large-scale nucleofections were performed by splitting cultures and conducting multiple parallel nucleofections.

Editing outcomes were measured four days post-nucleofection by flow cytometry or by amplicon sequencing (see below). Resuspension buffer and electroporation conditions were the following from each cell line: K562 in Buffer SF with FF-120, HEK293T in Buffer SF with DS-150, T cells in buffer P3 with EH-115, HCT116s in Buffer SE with EN-113, and HeLa cells in Buffer SE with CN-114.

### Genomic DNA Extraction (for Amplicon Sequencing)

Approximately 1e+05 cells were harvested and resuspended in 50uL of QuickExtract DNA extract solution (Lucigen). Reactions were incubated for 20 minutes at 65°C and 5 minutes at 95°C. Extractions were diluted 1:4 with dH2O, spun for 5 minutes at max speed in a microcentrifuge, and the supernatants retained for downstream analysis.

### PCR Amplification of Edited Regions

PCR reactions were generated from 2x Q5 master mix (NEB), primers [**Document S1**] at 500nM, and 5μL of genomic DNA (see above). Unless otherwise noted, PCR primers have a 5’ sequence tag (GCTCTTCCGATCT) that allows re-amplification for Illumina sequencing (amplify-on PCR). The thermocycler was set for one cycle of 98°C for 1 min, 35 cycles of 98°C for 10 sec, 63°C for 15 sec, 72°C for 60 sec, and one cycle of 72°C for 4 min, and held at 4°C. PCR amplicons were purified using SPRI beads, run on a 1.5% agarose gel to verify size and purity, and quantified by Qubit (Thermo Fisher, Waltham, MA).

### NGS Library Generation and Sequencing

Illumina adaptors and index sequences were added to 100ng of purified PCR amplicons by amplify-on PCR. Amplify-on was performed using 100ng of template DNA, 0.5 μM of forward/reverse primers, and 2x Q5 Master Mix (NEB). The thermocycler was set for one cycle of 98°C for 30 seconds, 16 cycles of 98°C for 10 sec, 62°C for 20 sec, 72°C for 30 sec, and one cycle of 72°C for 1 min, and held at 4°C. Each adaptor-conjugated amplicon was quantified by qubit, normalized, and pooled at equimolar amounts. Pooled samples were purified using SPRI beads. Library size and purity was verified by Bioanalyzer trace prior to sequencing on an Illumina MiSeq using reagent kit v3 (2×300bp).

### NGS Analysis of Amplicons

Samples were deep sequenced on an Illumina MiSeq at 300bp paired-end reads to a depth of at least 10,000 reads per sample. A customized version of CRISPResso ^38^ was used to analyze editing outcomes. Briefly, reads were adapter trimmed then joined before performing a global alignment between reads and the reference and donor sequences using NEEDLE ^39^. Rates of HDR are calculated as total number of reads that are successfully converted to the donor sequence (and have no insertions or deletions at the cutsite) divided by the total number of aligned reads. NHEJ rates are calculated as any reads where an insertion or deletion overlaps the cutsite or occurs within a six basepair window around the cutsite divided by the total number of aligned reads. SSTR/HDR rates were calculated at specific regions by counting total number of reads with flag occurrence divided by the number of aligned reads.

### Pooled Screen

Replicate cultures of K562 cells stably expressing a dCas9-KRAB construct and a cassette containing a BFP reporter and a guide RNA targeting a library of DNA repair factors (previously described^8^) were thawed, cultured, and puromycin treated. 10e+06 cells from each replicate were subcultured (UNZAP). 25e+06 cells from each replicate were harvested for nucleofection. Each nucleofection aliquot was spun down, washed in PBS, and resuspended in 825 μl of nucleofection buffer SF (Lonza, Basel, Switzerland). 275 μl of RNP editing mixture was added and mixed by pipetting. RNP for each replicate contained 2000 pmoles of sgRNA, 1,650 pmoles of Cas9, and 220 μg of plasmid pCR1075 donor DNA in Cas9 buffer (20 nM HEPES [pH 7.5]), 150 mM KCl, 1mM MgCl_2_, 10% glycerol, and 1 mM TCEP). Each replicate of the RNP cell slurry was split and nucleofected in parallel in a Lonza 96-well Shuttle nucleofector (code FF-120), re-pooled, and cultured for two (replicate 1) or three (replicate 2) days. Nucleofected replicates were sorted into GFP+ (GFP) and non-fluorescent (NON) populations on a Sony SH800S sorter. In parallel, an 85e+06 cell aliquot was harvested from each non-electroporated population the day of the sort (UNZAP), and an 85e+06 cell aliquot was harvested from each nucleofected cell library on the day of sorting (PRESORT). Harvested and sorted populations were spun down, rinsed in PBS, and frozen at − 80°C.

DNA from each cell population: PREZAP, UNZAP, PRESORT, GFP, and NON (non-treated) was purified using Machery-Nagel Blood Purification kits and the total amount of DNA quantified. One microgram of genomic DNA was amplified per Phusion HiFi PCR reaction using primers specific to the gRNA cassette as described ^40^. Up to 24 PCR reactions were set up for each cell population to obtain desired coverage of the cell library. The thermocycler was set for one cycle of 98°C for 30 sec, 25 cycles of 98°C for 15 sec, 56°C for 15 sec, and 72°C for 15 sec, and one cycle of 72°C for 10 min and held at 4°C. PCR reactions were pooled, purified using SPRI beads, and sized on an agarose gel. Amplified DNA from each cell population was normalized to input cell numbers, purified a second time using SPRI beads, and sequenced on a HiSeq2500 (Illumina).

### Pooled Screen Analysis

Data analysis was performed as described ^10^. Briefly, sequence reads were trimmed, aligned to DNA Repair Guide Sequences [**Document S1 GUIDES**] and quantified. Read counts for each gRNA were normalized and compared to the distribution of untargeted control guides to determine significance and log2 magnitude of change. The top three guide-level phenotypes were collapsed to produce gene-level phenotype score. Results for the GFPvPRE comparison are available in [**Document S2**].

### Pooled Screen Phenotype Comparison

The magnitude of gene-level phenotype scores (calculated above) varied between screens. To facilitate direct comparison of essential genes between HR and SSTR results, essential genes (phenotype scores < 0) were unity normalized against all essential gene phenotype scores for the originating screen, such that Z=(phenotype_score - min(dataset))/(max(dataset)-min(dataset)). The resulting normalized values ranged from 0 (strongest phenotype score) to 1 (weakest phenotype score). Z values were binned for display purposes: bin 1, 0.0–0.2; bin 2, 0.2–0.4; bin 3, 0.4–0.6; bin 4, 0.6–0.8; and bin 5, 0.8–1.0. FA and related genes from either screen (FAAP20, UHRF1, TIP60, POLQ, and LIG4) with a phenotype score > 0 were assigned to bin 5. Raw data from this figure is presented in [**Document S2**].

### Pooled Screen GO-term Comparison

Data from SSTR^8^ and HR (this manuscript) screens was filtered for statistical significance (p>0.05) and separated into two categories: genes involved in SSTR or genes involved in HR. The gene list for each category was compared to the starting guide pool [**Document S1 GUIDES**] using DAVID v6.8^13^. Default search categories were used.

### siRNA Transfection

Approximately 1e+05 suspension or adherent cells were reverse transfected into 24-well plates using RNAiMAX (Thermo Fisher). siRNA [**Document S1**] dosage was 40 nM unless otherwise indicated. Cells were siRNA treated for 48 hours, harvested, nucleofected, and recovered into media lacking siRNA. Verification of knockdown was performed at the time of nucleofection via qPCR. Cells were harvested for flow cytometry or amplicon sequencing 4 days post nucleofection.

### Drug Treatment

Acyriaflavin A, XL413, Aphidicolin, GW843682X and Ro3280 were sourced from Tocris. Hydroxyurea, A64 trifluoroacetate, GKT136901, SB220025 and UNC2170 trifluoroacetate were sourced from Sigma.

Approximately 1e+05 suspension or adherent cells were seeded post-nucleofection into 96-well plates in drugged media with the following concentrations: Acyriaflavin A at 5 μM, XL413 at 33 μM, Hydroxyurea at 2 mM, SRC7 at 1 μM, RS1 at 10 μM, Aphidicolin at 2 μg/mL, A64 trifluoroacetate at 1 μM, SB220025 at 0.5 μM, GKT136901 at 50 μM, GW843682X at 0.5 μM, Ro3280 at 100 nM, and UNC2170 trifluoroacetate at 150 μM. Cells were drug treated for 24 hours, harvested, washed, and recovered into fresh media. Cells were harvested for flow cytometry or amplicon sequencing 4 days post nucleofection.

### qPCR

Between 1e+05 and 2e+05 cells were harvested and RNA extracted using Qiagen RNeasy kits (Qiagen, Venlo, Netherlands). cDNA was produced from 1 μg of purified RNA using the iScript Reverse Transcription Supermix (Bio-Rad). qPCR reactions were performed using the SsoAdvanced Universal SYBR Green Supermix (Bio-Rad) in a total volume of 10 μl with primers at final concentrations of 500 nM. The thermocycler was set for 95°C for 2 mins and 40 cycles of 95°C for 2 sec and 55°C for 8 sec. Fold enrichment of the assayed genes over the control *ACT1B* and/or *GAPDH* loci were calculated using the 2^−ΔΔCt^ method essentially as described ^41^. Primer sequences can be found in **Document S1**.

### Western Blotting

Cells were harvested and washed with PBS. Cell were lysed in 1x RIPA buffer (EMD Millipore) for 10 min on ice. Samples were spun down at 14,000 × g for 15 min, and cleared protein lysates were transferred to a new tube. 20 μg of RIPA protein lysate was resolved on a TGX anyKD gel (Bio-Rad) and semi-dry transferred (TransBlot Turbo, Bio-Rad) to nitrocellulose membranes. Membranes were blocked in 5% milk, incubated with primary antibodies in 5% milk, incubated with secondary antibodies in 5% milk, and exposed on a LiCor Odyssey CLx (Li-Cor). Antibodies used were: FLAG (Sigma F1804, 1:1000), TOMM20 (Cell Signaling 42406, 1:1000), GAPDH (cell Signaling 97166, 1:5000), phospho-S53 MCM2 (Abcam ab109133, 1:1000), MCM2 (Abcam ab6153, 1:1000), 1:10,000 donkey anti-mouse IgG-IR800 (Li-Cor 925-32212), 1:10,000 donkey anti-mouse IgG-IR680 (Li-Cor 925-68022), 1:10,000 donkey anti-rabbit IgG-IR800 (Li-Cor 925-32213), 1:10,000 donkey anti-rabbit IgG-IR680 (Li-Cor 925-68023).

### T-Cell Experiments

Primary human T cells were isolated from two de-identified healthy human donors from residuals from leukoreduction chambers after Trima Apheresis (Blood Centers of the Pacific). Peripheral blood mononuclear cells (PBMCs) were isolated by Ficoll centrifugation using SepMate tubes (STEMCELL, per manufacturer's instructions), then T cells were further isolated from PBMCs by magnetic negative selection using an EasySep Human T Cell Isolation Kit (STEMCELL, per manufacturer’s instructions). Isolated T cells were cultured at 1 million cells/mL in XVivo15 medium (STEMCELL) with 5% Fetal Bovine Serum, 50 mM 2-mercaptoethanol, and 10 mM N-Acetyl L-Cystine, and stimulated for 2 days prior to electroporation with anti-human CD3/CD28 magnetic dynabeads (ThermoFisher) at a beads to cells concentration of 1:1, along with a cytokine cocktail of IL-2 at 200 U/mL (UCSF Pharmacy), IL-7 at 5 ng/mL (ThermoFisher), and IL-15 at 5 ng/mL (Life Tech). T cells were harvested from their culture vessels and magnetic anti-CD3/anti-CD28 dynabeads were removed by placing cells on an EasySep cell separation magnet for 3 minutes. Immediately prior to electroporation, de-beaded cells were centrifuged for 10 minutes at 90g, media aspirated, and resuspended in the Lonza electroporation buffer P3 using 20 μL buffer per one million cells.

RNPs were generated as described immediately prior to electroporation. Briefly, crRNA targeting the N-terminus of *RAB11A* (guide sequence GGUAGUCGUACUCGUCGUCG) and tracrRNAs were chemically synthesized (Edit-R, Dharmacon Horizon), and Cas9-NLS was recombinantly produced and purified (QB3 Macrolab). Lyophilized RNA was resuspended in 10 mM Tris-HCL (7.4 pH) with 150 mM KCl at a concentration of 160 μM, and stored in aliquots at −80C. crRNA and tracrRNA aliquots were thawed, mixed 1:1 by volume, and annealed by incubation at 37C for 30 min to form an 80 μM gRNA solution. This was further mixed 1:1 by volume with 40 μM Cas9-NLS protein to achieve a 2:1 molar ratio of gRNA:Cas9, with final RNP concentration of 20 μM. A long double-stranded HR template creating an N-terminal fusion protein of GFP and the *RAB11A* gene (Roth et al, Nature 2018) was generated as a PCR amplicon using KapaHiFi polymerase (Kapa Biosystems), purified by SPRI bead cleanup, and resuspended in water to 125 ng/uL as measured by light absorbance on a NanoDrop spectrophotometer (Thermo Fisher).

50 pmol of RNP and 0.25 μg of dsDonor were mixed for 5–10 minutes, then added to cells 3–5 minutes before electroporation. One million T-cells per well with RNP and dsDonor were electroporated using the Lonza 4D 96-well electroporation system with pulse code EH115, in biological replicate of n=3. Immediately post-electroporation, prewarmed media was added to rescue the cells then each electroporation condition was split into 5 wells of a 96-well U-bottom tissue culture plate. Electroporated cells were incubated at 1 million cells/mL in final volume 200uL media with IL-2 at 500 U/mL and increasing concentrations of XL413. After 24 hours, all cells were washed in PBS, and fresh media was added containing only IL-2 at 500 U/mL. Approximately 3 days post-electroporation, cells were collected by centrifugation at 300g, media discarded, and antibody stains were added (UCHT1-CD3-PE, OKT4-CD4-PE-Cy7, RPA-T8-APC (all from BioLegend), and GhostDye780 (Tonbo)) for 20 minutes. Cells were washed, resuspended in PBS+ 1% serum (120uL per well), and then an equal volume (80uL) of each well was sampled using an Attune NxT Focusing Flow Cytometer with Autosampler attachment (ThermoFisher).

### FUCCI Cell Line Generation

K562 cells were electroporated with 40 μg pML143 donor vector containing EF1α-mAG1-geminin(1-110)-P2A-mKO2-hCdt1(30-120) flanked by homology arms to AAVS1/PPP1R12C locus and 5μg each of AAVS1L and AAVS1R TALENs. Targeted cells were sorted by FACS into 96 well plates to isolate single clones. Clonal cell lines were assayed by live cell imaging and flow cytometry to verify correct correlation with fluorescence and cell cycle progression.

### Cell Cycle Assay/DNA Quantification Assay

K562 FUCCI cells were grown in complete RPMI containing Aphidicolin (2 μg/mL), Hydroxyurea (2 mM) or XL413 (33 μM). Cells were harvested at indicated time points and subjected to flow cytometry. Cell cycle status was determined gating for mAG1^+^/mKO2^−^ (S/G2/M), mKO2^+^/ mAG1^−^ (G1), mAG1^+^/ mKO2^+^ (early S) and mAG1^−^/mKO2^−^ (early G1) cell populations. DNA content of the cells was determined using Hoechst33342 (Thermo Fisher) DNA dye. Hoechst33342 was added to cells in culture medium (final concentration 1 μg/mL) for 30 mins at 37 °C before cells were subjected to flow cytometry.

### Data Availability

Data from CRISPRi screens will be publicly available at [**Database Location**].

